# An Empirical Energy Landscape Reveals Mechanism of Proteasome in Polypeptide Translocation

**DOI:** 10.1101/2021.06.21.449244

**Authors:** Rui Fang, Jason Hon, Mengying Zhou, Ying Lu

## Abstract

The ring-like ATPase complexes in the AAA+ family perform diverse cellular functions that require coordination between the conformational transitions of their individual ATPase subunits^1,2^. How the energy from ATP hydrolysis is captured to perform mechanical work by these coordinated movements is not known. In this study, we developed a novel approach for delineating the nucleotide-dependent free-energy landscape (FEL) of the proteasome’s heterohexameric ATPase complex based on complementary structural and kinetic measurements. We used the FEL to simulate the dynamics of the proteasome and quantitatively evaluated the predicted structural and kinetic properties. The FEL model predictions were widely consistent with experimental observations in this and previous studies and suggested novel features of the mechanism of proteasomal ATPase. We find that the cooperative movements of the ATPase subunits result from the design of the ATPase hexamer entailing a unique free-energy minimum for each nucleotide-binding state. ATP hydrolysis dictates the direction of substrate translocation by triggering an energy-dissipating conformational transition of the ATPase complex.

## Introduction

The ring-shaped oligomeric ATPases control key biological processes including protein folding, transcription, DNA replication, cellular cargo transport and protein turnover^1–3^. The 26S proteasome is an ATP-dependent protein degradation machine in the AAA+ (<u>A</u>TPases <u>a</u>ssociated with diverse cellular <u>a</u>ctivities) family of ATPases^4^. The proteasome holoenzyme consists of a barrel-shaped 20S core particle (CP) capped by 19S regulatory particles (RP) on one or both ends^5^. Each RP features a nine-member Lid subcomplex and a heterohexameric ring of AAA+ ATPases assembled from six distinct gene products (RPT1-RPT6) sharing 85% sequence identity. These ATPases are responsible for mechanically unfolding substrates using the energy from ATP hydrolysis and translocating them into the CP for proteolysis (Fig. 1A).

**Figure 1.**
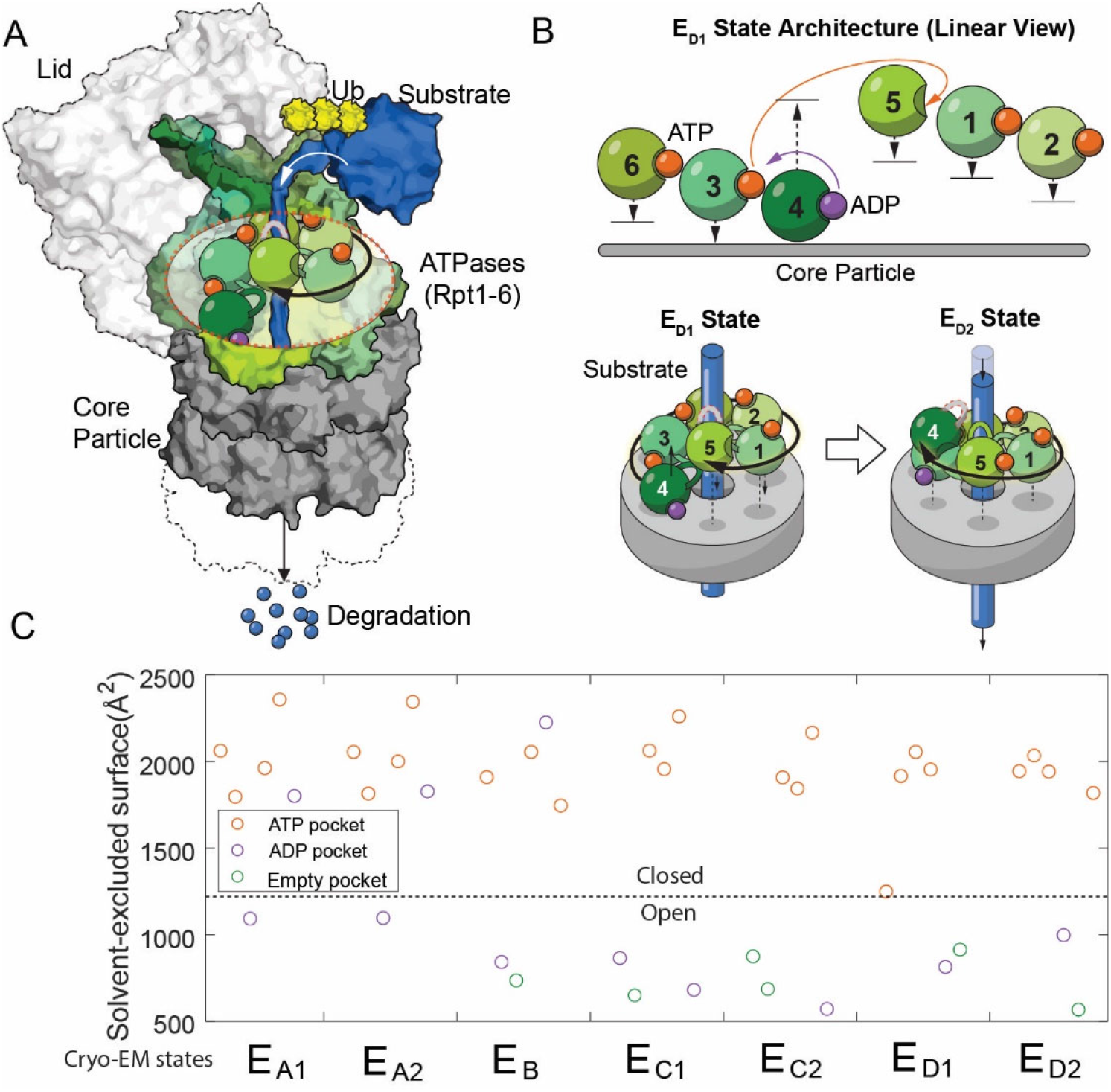
The architectures of the proteasomal ATPase complex and its interaction with substrate. **A.** A schematic showing a half 26S proteasome engaged with an unfolded substrate through the PL1s (color loops) on the ATPase subunits with bound nucleotides (color blobs). The disengaged PL1 is marked in grey. A black arrow suggests the staircase architecture of the ATPases. **B.** Upper panel: a linear view of the architecture of the ATPase complex in a translocating state E_D1_. Each ATPase subunit (Rpt1~Rpt6) is shifted vertically according to the position of its PL1 relative to the core particle. An open interface is suggested by a large gap between subunits. Arrows indicate the displacements of the PL1s and the change of nucleotides in the E_D1_-to-E_D2_ transition. See Fig. S1 for a complete list of the identified states of the proteasome. The lower panel illustrates the mechanism of substrate translocation coupled with the ATPase rearrangement in the E_D1_-to-E_D2_ transition: the PL1 on Rpt4 disengages from substrate and moves to the distal registry of the staircase. This change is accompanied by an axial movement of the PL1s on Rpt1/2/6/3 that still interact with the substrate towards the core particle to bring about axial stepping and translocation of approximate 2xAAs. **C.** The solvent-excluded surface area at the interfaces of the ATPase domains in the seven substrate-engaged proteasome structures, colored according to the nucleotide at each binding pocket. The dashed line separates the closed and open interfaces, as defined here.

Processive translocation and degradation of protein substrates is critical for the biological functions of the ubiquitin-proteasome system^4,6^. This process requires highly-coordinated conformational changes of the ATPase units on proteasome. To reveal the structural mechanism of proteasomal degradation, previous studies using cryo-electron microscopy (cryo-EM) of the substrate-engaged proteasome captured seven states, named E_A1_-E_D2_, with E_D1_ & E_D2_ hypothesized involved in substrate translocation (Fig. S1)^7,8^. An unfolded substrate primarily interacts with aromatic residues on the pore-1 loop (PL1) of each ATPase. These short structured loops form a right-handed helical “staircase” delineating the interior of the translocation channel. Changes in bound nucleotides at the ATPase interfaces are associated with rearrangements of the architecture of the PL1 staircase and the proteasome-substrate interaction, as a result of both translational and pivot movements of these ATPases (Fig. 1B and fig. S1)^7,8^. The PL1s that are disengaged from substrate interaction usually occupy distal, or top, positions in the staircase away from the CP (Fig. 1A). When certain PL1s disengage and move to the top, this is accompanied by a movement of substrate-engaged PL1s in the proximal direction–towards the CP. This conformational rearrangement may provide the power stroke to promote axial translocation of substrate (Fig. 1B)^7,8^.

The activities of proteasome are broadly driven by its structural dynamics which remains largely elusive. Based on these cryo-EM structures, we previously proposed that the conformation and nucleotide-binding status of the ATPase hexamer may cycle consecutively through six different states with rotational equivalence, thus driving processive substrate translocation^7,9^. A similar model of sequential transitions was proposed in an independent study of yeast proteasome structures^8^. Direct experimental validation of this sequential-transition model has not yet been reported, and only two out of the six conformations proposed in the sequential-transition model were identified in these structural studies^7,8^. In spite of these structural insights, we still do not know how the chemical energy from ATP hydrolysis is harvested to drive the ordered transition of proteasomal conformations, and in particular how individual ATPase subunits coordinate in their chemical and mechanical cycles to achieve this.

In addition, there are experimental findings that appear inconsistent with the predictions of a strict sequential-transition model. For example, mutation of the Walker-A (WA) or Walker-B (WB) motif on an ATPase, which impedes its nucleotide-binding or hydrolysis activity, would inactivate the proteasome by blocking the transition sequence. However, such mutations on some of these ATPases are in fact well tolerated in yeast^10–12^. Mutating other functional motifs on different proteasomal ATPases also has distinct effects on protein degradation, despite their high levels of similarity in sequence and structure, leading to the hypothesis that the six ATPases may play nonequivalent roles in the proteasome activities^10,13,14^. These functional properties of the proteasome are not interpreted by the previous models and alternative models that are consistent with both structural and functional observations have not been suggested.

To identify the mechanism of proteasomal ATPases in substrate translocation, we simulated the conformation dynamics of the ATPase complex by constructing a physical model based on the nucleotide-dependent free-energy landscape (FEL) of the ATPase complex. To obtain the FEL, We first performed comparative analysis on known proteasome structures to identify the primary degrees of freedom (DOF) of proteasome’s conformational changes, and then parameterized the free energy surface, or potential of mean force, based on the mode of nucleotide-ATPase interactions, and experimentally determined the FEL parameters. Simulation of the FEL model generates testable predictions that are separate from the results for model construction. We then experimentally evaluated these predictions in a wide range of conditions, as well as comparing them with published results, and found that most of these predictions were congruent with the experimental measurements in this and previous studies.

Our work introduces a new method for studying the dynamics of a complex protein machine such as the 26S proteasome, and demonstrates that this FEL approach not only provides a coherent explanation for a variety of structural and kinetic observations but also reveals the underlying mechanism by which the AAA+ hexamer on the proteasome operates in driving substrate translocation.

## Results

### I. Structure-based construction of the FEL of the proteasomal ATPase complex

In this work we elect a combination of the ATPase complex’s conformation and the nucleotide distribution in the six ATPase pockets to specify a “state” of the proteasome, for consistency with nomenclature in structural studies. A description of the FEL of proteasome is represented by the potential of mean force of a specific proteasome population measured as a bivariate function of its conformational coordinates and the nucleotide distribution^15,16^.

The primary DOF of proteasome’s conformational changes identified by comparing proteasome structures define the conformational coordinates of the FEL. We designate each conformation in the FEL by whether the interfaces of the six ATPase domains on proteasome are open or closed, based on the observation that the solvent-excluded surface area (SESA) of these interfaces mostly adopts binary values in these cryo-EM structures (Fig. 1C and fig. S2). A large SESA is associated with closed interfaces that appear to arrange the PL1s on neighboring ATPases into a staircase. In contrast, a smaller SESA suggests an open interface and the relative geometry of the neighboring ATPases tends to vary (Fig. S1). In the FEL, we constrain the total number of open interfaces in each hexamer conformation to be either 2 or 3, as observed in the cryo-EM structures except for E_A_ states which involve only one open interface (Fig. S1) (see Discussion). This defines 30 conformations, after excluding 5 with ambiguity in assigning proteasome-substrate interaction due to symmetry (Table S1; methods 9.2). States in the FEL are also differentiated by the nucleotide distribution in these ATPases. Each nucleotide pocket may be occupied by ATP, ADP, ATP-γS(a slowly-hydrolyzing ATP analog) or no nucleotide. The total number of distinct states in the FEL model is therefore 30^x^4^6^=122,880. To accelerate the simulation, we ignore the transient ADP•Pi state due to the very weak affinity between free phosphate and the proteasome (Fig. S3), and no such state has been identified so far in proteasome structures.

The molecular details of the nucleotide-ATPase interactions suggest a strategy to parameterize the free energy of each conformation as a function of its nucleotide status. Each nucleotide-ATPase interaction on the proteasome involves several conserved elements: the WA, WB, sensor I and sensor II motifs from the *cis* ATPase and the arginine fingers from the *trans* ATPase (Fig. 2A)^17^. The arginine fingers are the major components interacting with the γ- and β-phosphate groups on ATP at a closed interface. We found that the *cis* elements exhibited rather minor rearrangements between different proteasome states (RMSD ~ 0.58Å) (Fig. S4). We therefore parameterized the total free energy of the ATPase hexamer as a sum of the contributions from each individual ATPase interface. Each interface contribution is subdivided into three components: the basal energy *E_b_*, which derives from direct interactions between adjacent ATPases at a closed interface; the pocket energy *E_p_* from the nucleotide-*cis*-element interactions, which are similar for ADP and ATP; and the bridge energy *E_Br_*, which differentiates ADP from ATP and originates from the engagement of arginine fingers and other *trans* elements with the γ- and β-phosphate on the nucleotide at a closed interface (Fig. 2B). The *cis* pockets of disengaged ATPases exhibit either low or no nucleotide density in cryo-EM maps, and are associated with slightly rearranged *cis* motifs (Fig. S1 and fig. S4)^7^. We therefore assign a separate pocket energy 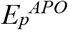 for disengaged ATPases to reflect their low affinity for nucleotides. The chemical energy in ATP is excluded from the free energy calculation, since it does not explicitly contribute to the simulation of proteasome dynamics (see section II).

**Figure 2.**
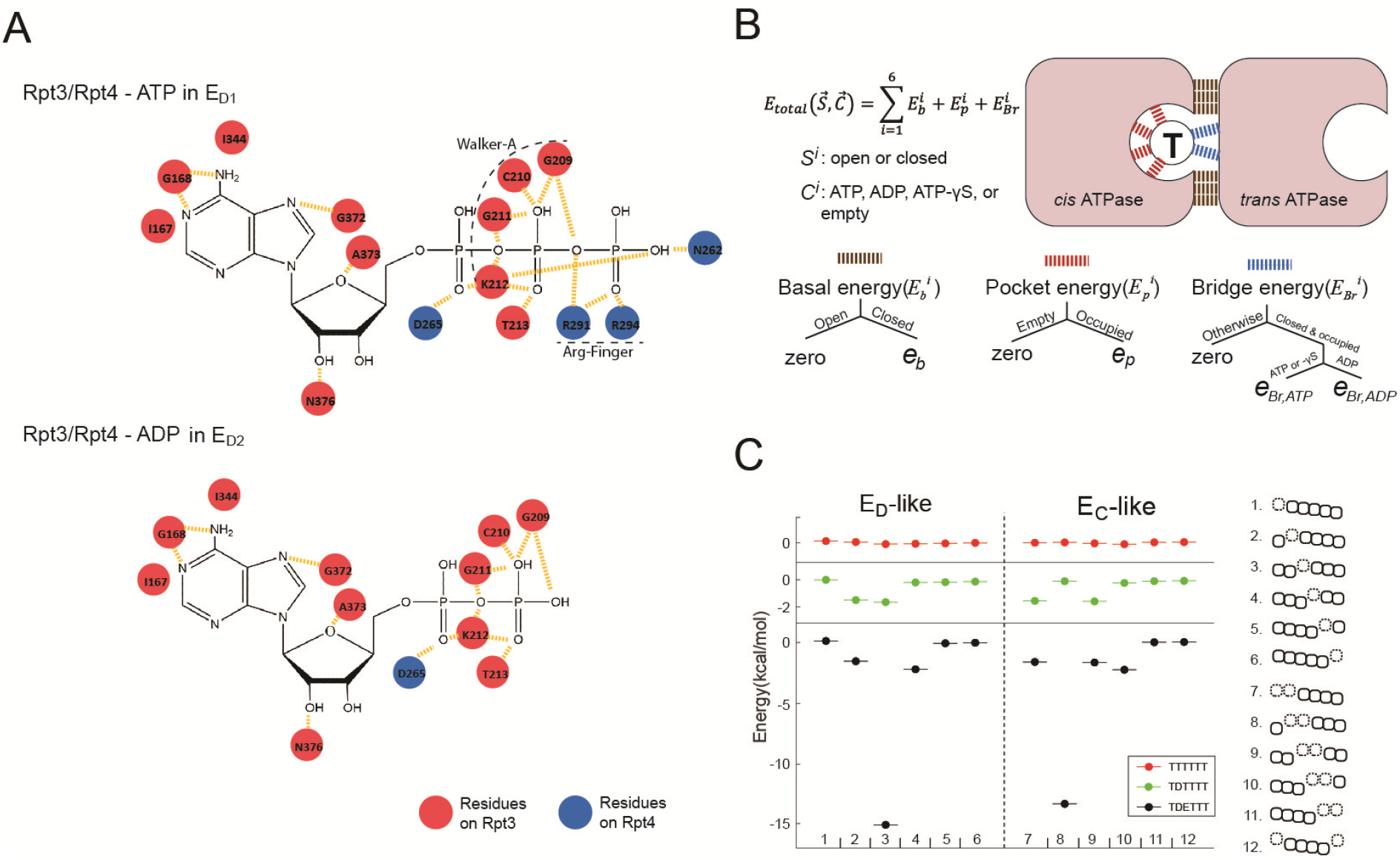
Parameterization of the nucleotide-dependent free-energy landscape. **A.** The interaction map of the residues on Rpt4 and Rpt3 with the bound ATP or ADP in the E_D1_ or E_D2_ states. Red: *cis*-interacting residues on Rpt3. Blue: *trans*-interacting residues on Rpt4. **B.** A schematic showing the parameterization strategy for the free energy (*E_total_*) of the ATPase hexamer and valuation of the parameters. The molecular interactions underpinning the three energy terms are marked by different colors. “Open/Closed” refers to the status of an ATPase interface *S^i^*. “Empty/Occupied/ADP/ATP/ATP-γS” refers to the status of the nucleotide-binding pocket in the *cis* ATPase *C^i^*. See methods 9.3 for a detailed explanation. **C.** The FEL on the E_D_-like and E_C_-like conformations of the ATPase complex in three representative nucleotide-binding statuses. T/D/E: ATP/ADP/empty. A sketch for the ATPase architecture of each conformation is listed on the right. The ATPases Rpt6/3/4/5/1/2 are represented by squares from left to right; dashed squares = disengaged ATPases that are flanked by open interfaces (gap between squares); the axial position of a PL1 is indicated by the vertical shift of the corresponding ATPase square.

These energy parameters can in principle vary for the different ATPase subunits Rpt1~Rpt6. In the analysis below, we made the simplifying assumption that all six ATPases share an identical set of parameters. This assumption does not contradict the experimental finding of *functional* disparity among the six ATPases. In section IV, we analyzed the contribution of the proteasome Lid-ATPase interaction to the symmetry breaking among the ATPases.

### II. Evaluating the FEL model parameters

We determined all nine parameters in the FEL model from the measurements in this study and previous work (Methods 9.3 & 9.4). A critical parameter is the difference in the bridge energies (*E_Br_*) between ATP and ADP interfaces, which we find is key for determining the directionality of translocation. This parameter was measured using a single-molecule binding assay, based on the relationship between the dissociation constant (*K_d_*) of nucleotides and these energy terms. In the FEL framework, we identified three affinity groups, each with a distinct *K_d_*, and the group designation varies with the status of the *cis* ATPase and the interface (Fig. 3A). We used a very low concentration (200nM) of Alexa647-conjugated ATP to limit the interaction to the strongest binding pockets (group 1 in Fig. 3A) on the human 26S proteasome which was immobilized on a passivated coverslip, and observed its interaction with ATP by TIRF (Total Internal Reflection Fluorescence) microscopy. To enhance the specificity of single-molecule detection, the proteasome was fluorescently labeled on the 19S particle using a SNAP tag for colocalization analysis with Alexa647-ATP. We varied the concentrations of unlabeled nucleotides to compete with Alexa647-ATP in a steady-state measurement, to circumvent the possibility that a conjugated fluorophore may affect the nucleotide-ATPase affinity in direct measurements (Fig. 3B). ATP-γS was employed to minimize nucleotide hydrolysis. This analysis yielded the following ratio of the inhibitor constant (*K_i_*) for ATP: ADP: ATP-γS = 1: 7.9 (±2.0): 0.5 (±0.15) (at 90% confidence interval), giving *e*_*Br*, ATP_ - *e*_*Br*, ADP_ =−1.6(±0.4) kcal/mol.

**Figure 3.**
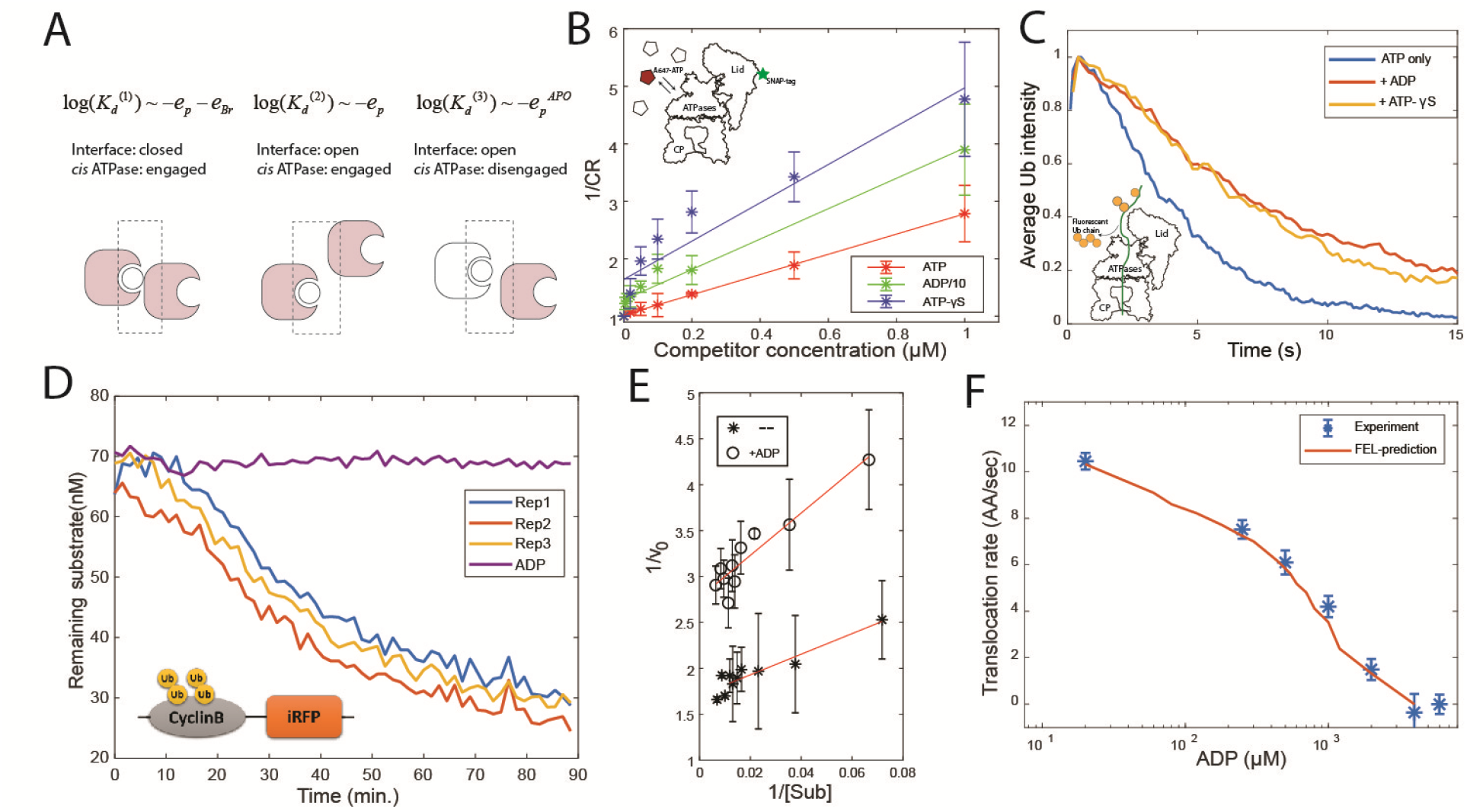
Evaluation of the FEL model parameters. **A.** A schematic showing three categories of nucleotide pockets with their corresponding dissociation constants *K_d_*. **B.** Single-molecule nucleotide-proteasome interaction assay. 200nM Alexa647-ATP (red pentagon) was mixed ATP, ADP or ATP-γS (white pentagons) as a competitor at different concentrations, incubated with surface-immobilized 26S proteasome which was labeled with a SNAP-tag dye. The degree of Alexa647-ATP-proteasome colocalization in a steady state was measured using a TIRF microscope. The colocalization ratio (CR) after normalization by the competitor-free condition was inversely regressed on the competitor concentration to obtain the relative inhibitor constant *K_i_* (methods 8). ADP concentration is divided by 10 for presentation. The inset illustrates the experimental design. Error bars represent the standard deviation of three replicas. **C.** Singlemolecule translocation assay. N-terminal cyclinB was conjugated with Dylight550-labeled ubiquitins (yellow disks) on lysines(l8, 36, 64) and was incubated with surface-bound 26S proteasome in a buffer containing 0.5mM ATP or with extra 0.8mM ADP or with extra 40μM ATP-γS. About 100 single-molecule traces exhibiting processive deubiquitylation in each condition were aligned by the time of substrate-proteasome encounter (t=0). The average fluorescent-ubiquitin intensity on a substrate molecule at each time point was calculated. The rate of translocation was calculated from the initial slope of the traces. The inset illustrates the experimental design. **D.** Representative traces of the degradation of ubiquitylated cyclinB-iRFP by 1.5nM purified 26S proteasome with ATP (rep1~3) or ADP in the buffer. **E.** Lineweaver-Burk plot of the initial degradation rate (v_0_) at varying concentrations of cyclinB-iRFP (Sub) either with 0.5mM ATP (--) or with extra 0.8mM ADP (+ADP). **F.** The translocation rate of cyclinB-iRFP was measured using the fluorescent degradation assay with 0.5mM ATP and various concentrations of ADP-Mg^2+^. Error bars represent the standard deviation of 15 measurements. The red curve shows the prediction by the FEL model.

After determining the FEL parameters, we explored its basic features. For the twelve E_D_-like and E_C_-like ATPase-hexamer conformations that are characterized by either a single or two adjacent disengaged ATPase units, uniform ATP binding generates a flat FEL. The presence of ADP in a single pocket breaks the symmetry and lowers the relative free energy of a subset of conformations by 1.6 kcal/mol. The further addition of an empty pocket next to the ADP pocket introduces a well-separated energy-minimum conformation in which the empty and ADP *cis* pockets are found in the disengaged ATPase and its counterclockwise neighbor (Fig. 2C). This configuration was identical to those observed in the E_D1_ and E_D2_ states of proteasome, as well as in related AAA+ ATPases (Fig. S1)^7,18,19^.

We simulated the dynamics of the ATPase complex as stochastic transitions in a discrete 6+6 dimensional space (Fig. S5). The transitions between any conformations were described as a simple bi-state process with a rate constant determined by the Arrhenius equation. The ATP cycle at each pocket proceeds independently, as described by the basic chemical rate equations. The off-rate of a nucleotide is calculated from its *K_d_*, with the on-rate set at a constant (Fig. 3A; Methods 9.3 & 9.4). The detailed process of a conformational change is not considered explicitly (i.e. as “adiabatic”) in that the change of nucleotide distribution alters the FEL and causes repopulation of the conformational space of proteasome. We do not include any assumption on the coordination of the nucleotide cycles or the coupling of the nucleotide and conformational changes.

For the simulation of ATPase dynamics to generate testable predictions, we introduced a substrate peptide that is mechanically coupled with the PL1 rearrangement through the proposed “hand-over-hand” mechanism (Fig. 1B)^7,8^. The unit step of translocation is defined by the axial separation of PL1s, corresponding to approximately two amino acids (AA) in the substrate polypeptide (Fig. 1B). The force coupling with a translocating substrate may affect the rates of ATPase conformational changes. We represent this effect as a titling of the FEL and simplified by ignoring the stochastic and sequence-dependent variations of these forces and introduced two constant values to capture the average effect in the forward and backward processes (Methods 9.4).

To experimentally determine these two force parameters and another parameter that defines the activation energy barrier, we employed a quantitative degradation assay based on the decay of a fluorescent reporter of the ubiquitylated cyclinB N-terminus fused with an infrared fluorescent protein iRFP (Fig. 3D)^20^. The N-terminal cyclinB is an unstructured protein that is efficiently degraded by the proteasome starting from its N-terminus^21,22^. Our analysis shows that degradation of cyclinB-iRFP follows Michaelis-Menten-like kinetics with *K_M_* = 5.6nM (Fig. 3E). We performed each measurement at substrate concentrations much higher than the *K_M_* to maximize the signal-to-noise ratio, and tested five substrate concentrations, each with three replicas, to verify the saturation condition. It took on average 55 seconds (i.e. the turnover time) for the proteasome to degrade a 46kDa cyclinB-iRFP molecule in a process that could be rate-limited by several steps including substrate commitment, unfolding and translocation^4^.

To determine the actual rate-limiting step, we engineered a mutant of cyclinB containing only three lysine residues, and conjugated fluorescent ubiquitin chains onto these lysines^23^. We examined this substrate using a single-molecule translocation assay developed previously (Fig. 3C)^23,24^. The processive translocation of a substrate coincides with the stepwise removal of entire ubiquitin chains by the deubiquitinase Rpn11 on the proteasome, which was detected by TIRF to measure the ubiquitin copies on a substrate molecule (Fig. S6). The decay rate of fluorescent ubiquitins on three-lysine cyclinB suggests a translocation rate of 10.5 (±0.8) AA/sec (Methods 7).

Given this measurement, translocating a cyclinB-iRFP peptide containing 445 AAs would take at least 44 seconds, or 80% of the 55-second total turnover time. This suggests that translocation is the limiting step of the entire degradation process for this substrate. Therefore, we measure the degradation rate of cyclinB-iRFP as an approximation for the rate of its translocation which is directly coupled to the ATPase conformational transitions. Degradation of these substrates is unlikely to be limited by deubiquitylation because substrates with multiple Ub chains are degraded faster than the same substrate with fewer chains^23^. This reporter assay also gives consistent results with direct single-molecule measurements under perturbed conditions, such as in the presence of ADP or ATP-γS (Fig. 3C). We next determined the translocation rate of cyclinB-iRFP in the presence of 500μM ATP and varying concentrations of ADP, and estimated the three translocation-related parameters using these data (Fig. 3F and methods 9.4).

### III. Examining the FEL-predicted kinetics of substrate degradation

To evaluate whether the simulation reflects the actual dynamics of the ATPase complex, we seek to examine the model predictions that are independent of the information for model construction and summarize the main results in Fig. S7. We first compared the simulated translocation rates of cyclinB-iRFP at different ATP concentrations with experimental measurements (Fig. 4A). Interestingly, the translocation-rate curve is non-monotonic and peaks around the physiological concentration of ATP (Fig. 4B). The FEL model accurately captures the quantitative features in both the up and down phases of the rate curve, each yielding a different insight. The EC_50_ value for ATP in the up phase is 45μM, far from the *K_d_* value of ATP or ADP at any binding pocket (Fig. 3A: ~100nM for group 1, ~3μM for group 2, ~2mM for group 3). Guided by the FEL model we derived the expression of this EC_50_ value as the ratio between the total rate of ATP hydrolysis and the on-rate of ATP (Methods 10). Inhibition of translocation at high ATP concentrations can also be rationalized by the FEL model. This inhibition is not due to extra Mg^2+^, proteasome disassembly, degradation-independent iRFP inactivation, or a general slowdown of ATP hydrolysis (Fig. S8 to S11). To achieve directional translocation of a substrate, there requires a unique energy-minimum ATPase conformation that varies with nucleotide-binding status and thus drives an ordered conformational change, leading to translocation. High ATP concentrations bias the proteasome towards all-ATP states and a flat FEL, resulting in a loss of the directionality of translocation (Fig. 2C, 4A). In contrast, a sequential-transition model would predict a monotonic increase of translocation rate at increasing ATP concentrations, inconsistent with the observation. (Methods 11).

**Figure 4.**
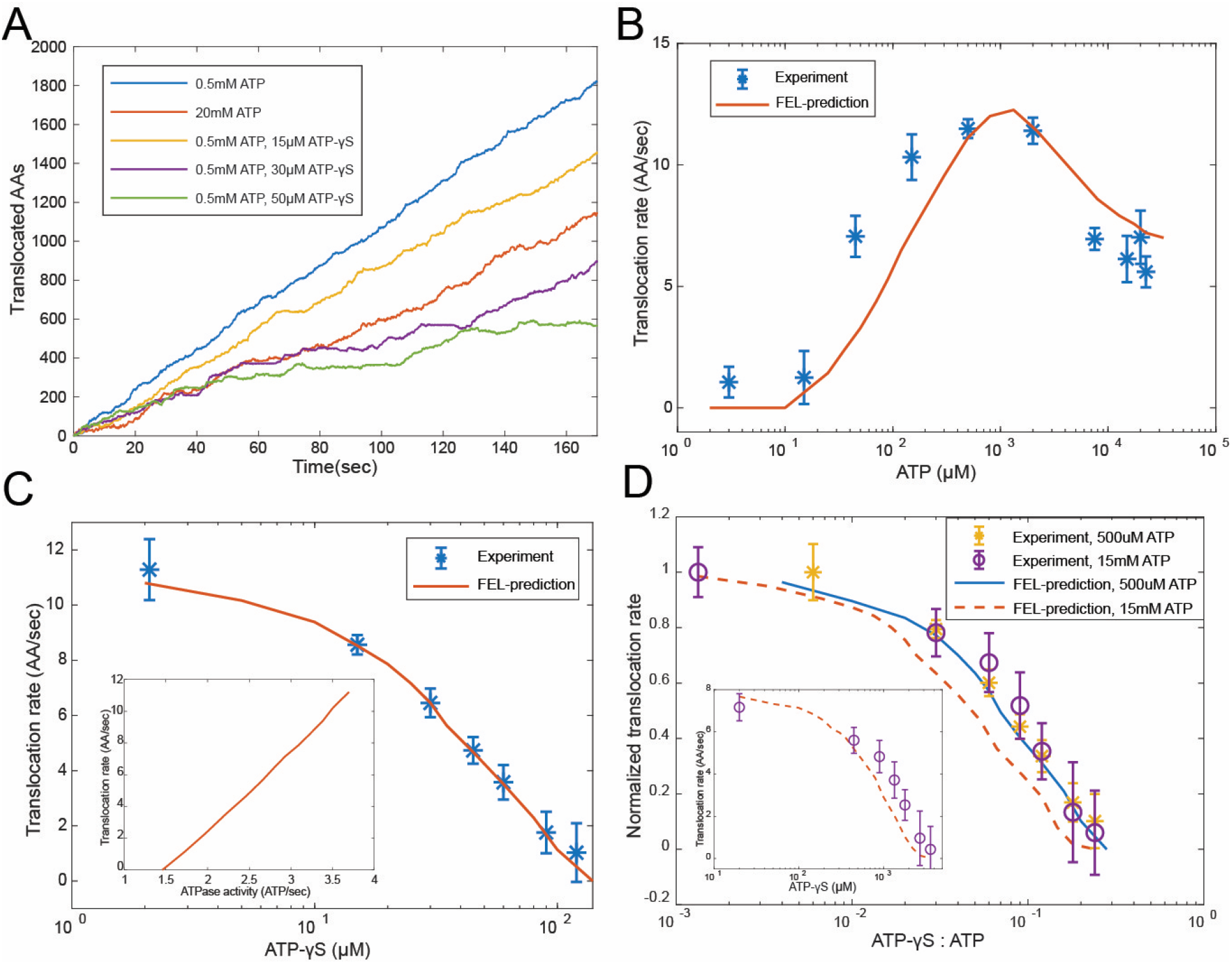
Evaluating the FEL-predicted degradation kinetics. **A.** Examples of simulated kinetics of translocation on individual proteasome particles under indicated nucleotide conditions. **B.** The translocation rate of cyclinB-iRFP measured at various concentrations of ATP-Mg^2+^, in comparison with the FEL-model prediction (red curve) based on the determined parameters. **C.** Same as in **B**, but measured with 0.5mM ATP and various concentrations of ATP-γS. Inset: translocation rate vs. ATP-hydrolysis activity as predicted by the FEL model at 0.5mM ATP and different concentrations of ATP-γS. **D.** Same as in **B**, but measured with l5mM ATP and various concentrations of ATP-γS. X-axis is the ratio between ATP-γS and ATP concentrations; Y-axis is the normalized translocation rate. The result in **C** is overlaid as a comparison. Inset: the absolute translocation rate vs. ATP-γS concentrations.

The FEL-predicted translocation kinetics are also consistent with the results of competition experiments using ATP-γS. In the simulation, ATP-γS introduces pauses in substrate translocation, longer at higher concentrations, which is reminiscent of the ATP-γS-induced translocation pauses in a single-molecule force measurement on ClpXP, a proteasome-like ATPase in prokaryotes (Fig. 4A)^25^. Here, 36μM ATP-γS inhibited the translocation rate by 50% in the presence of 500μM ATP (Fig. 4C), despite the fact that the difference in the apparent *K_i_* values between ATP and ATP-γS is only 2-fold (Fig. 3B). This 36μM-IC_50_ can be explained by considering other steps, such as ATP hydrolysis, in the FEL model. To test whether the FEL prediction is still valid at high ATP concentrations, we performed an ATP-γS competition experiment at a level where the high-ATP-inhibition effect is apparent (15mM ATP). Despite an overall reduction in translocation and a dramatic shift in IC_50_, the FEL-predicted rates still closely match the experimental results (Fig. 4D). The FEL model also predicts that the translocation rate should linearly depend on the rate of ATP hydrolysis at varying ATP-γS concentrations (Fig. 4C), consistent with a previous observation^26^.

For structurally-stable substrates, unfolding may be a limiting step in degradation by the proteasome. To test the FEL model in the context of such substrates, we created a fluorescent reporter by inserting a dihydrofolate reductase(DHFR) domain from *E. coli* between cyclinB and iRFP. We found that adding the DHFR ligand folic acid led to a dose-dependent stabilization of the ubiquitylated cyclinB-DHFR-iRFP in the presence of the proteasome, while the degradation of the original cyclinB-iRFP was unaffected (Fig. S12). In the presence of folic acid, cyclinB-DHFR-iRFP degradation is still complete, or processive (Fig. S13). The IC_50_ value for folic acid in inhibiting the degradation is 800μM, much higher than the 1μM dissociation constant between DHFR and folic acid^27^. This is likely because unfolding of the DHFR domain by the ATPase’s actions primarily occurs when DHFR is transiently unliganded, consistent with the linear relationship between folic acid concentration and the inverse of the degradation rate (Fig. S14 and methods 12). The FEL model suggests that folic acid lowers the overall efficiency of ATP utilization in degrading DHFR-containing substrates as reported previously (Fig. 5A)^26^. We tested the degradation rates of ubiquitylated cyclinB-DHFR-iRFP with 800μM folic acid in an ATP-titration experiment, and found that folic acid did not affect the EC_50_ value of ATP in the up-phase of the rate curve, although it lowered the peak degradation rate. However, folic acid significantly reduced the degree of degradation inhibition at high ATP concentrations (Fig. 5A, B). This is likely because the ATPase activity unfolding the substrate is less affected by high ATP concentrations than is translocation (Fig. S11, S15). To predict the degradation rate of cyclinB-DHFR-iRFP, we introduced one additional parameter to describe the unfolding rate of unliganded DHFR by proteasome in the FEL model. These qualitative features of the predicted degradation-rate curve in the ATP-titration experiment are insensitive to the choice of this parameter (Methods 12).

**Figure 5.**
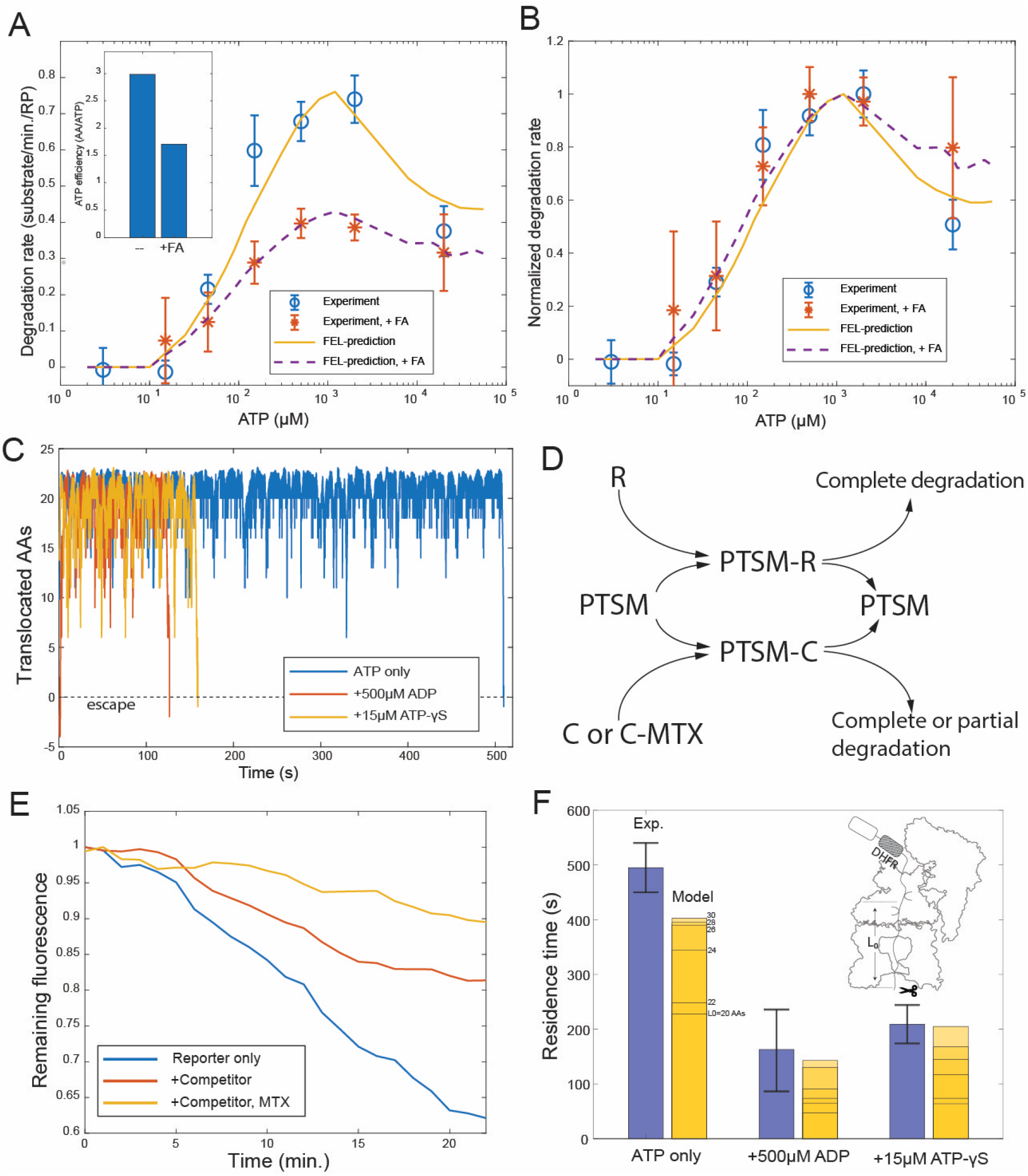
Evaluating the FEL predictions for structurally-stable substrates. **A.** The initial degradation rate of ubiquitylated cyclinB-DHFR-iRFP by purified 26S proteasome with or without 0.8mM folic acid (FA), overlaid with the FEL model prediction. The inset shows the effective length of substrate peptide degraded by consuming one ATP molecule, as predicted by the FEL model. The normalized degradation rate is shown in **B.** Error bars represent the standard deviation of 15 measurements. **C.** Examples of simulated translocation kinetics when the ATPases encounter an unfolding-resistant domain at t=0, assuming substrate escape occurs at Y=0. **D.** Schematic showing the reactions in the competition assay for determining the residence time of an unfolding-resistant substrate on proteasome. R: cyclinB-iRFP reporter; C: cyclinB-DHFR-iRFP^dark^ competitor. **E.** Representative traces of the competition assay in D. **F.** Experimental values of the residence time of cyclinB-DHFR(MTX)-iRFP under indicated nucleotide conditions, in comparison with FEL model predictions with the peptide track length L_0_ from 20 to 30AAs. Error bars represent the standard deviation of 5 measurements.

Molecular machines may occasionally run backwards due to the stochasticity of single-molecular dynamics. Simulation by the FEL model suggests that substrates with an unfolding-resistant domain would not stably engage with the proteasome but escape at a rate determined by the backward kinetics (Fig. 5C). A stable ligand, such as methotrexate, can inhibit the degradation of cyclinB-DHFR-iRFP at a low concentration by preventing the unfolding of DHFR, resulting in partial cleavage of the substrate (Fig. S12)^28^. In a competition assay, degradation of the reporter cyclinB-iRFP was reduced by a nonfluorescent competitor cyclinB-DHFR-iRFP^dark^ (Fig. 5D). And this effect was exacerbated in the presence methotrexate (Fig. 5E). We measured the turnover time of the stable DHFR substrate in this assay and found the value comparable with the model prediction which was calculated as a first-passage time on a 20~30AA peptide track measured from the PL1s to the proteolysis sites in the CP (Fig. 5F) (Methods 9.5 & 14). Addition of ATP-γS or ADP in the simulation reduces the first-passage time and facilitates the escape of stable substrates (Fig. 5C). This trend is also consistent with the measured turnover time in the presence of ATP-γS or ADP (Fig. 5F).

In summary, we found that predictions from our FEL approach using nine empirically-determined parameters closely matched new experimental observations at nucleotide concentrations ranging across three orders of magnitude, in both the forward and the backward processes. The overall consistency with experiments is not sensitive to parameter uncertainty at a typical value of ~30% (Fig. S16).

### IV. Organization of proteasomal conformations in the dynamical space

We next apply the FEL-based simulation to examine the global features of the conformational transitions of the proteasome. This analysis provides critical insights into the cooperative mechanism of the ATPase units and interprets the effects of ATPase mutations. We calculated the steady-state distribution of the proteasome at each conformation and the frequency of every conformational transition in simulating the FEL model for the translocation process. The FEL model is built on fundamental physical and chemical processes and does not *a priori* specify the occupancy of any conformation, the transition pathway between conformations, or how the ATPases cooperate. Interestingly, in the simulation we found that the ATPase conformations self-organized into a transition network, with the E_D_-like and E_C_-like conformations predominating, as in the cryo-EM analysis (Fig. 6A)^7^. Although the main transition pathway among EDs resembles the previous sequential model, several side transitions, or branches, can be identified, which are key to understanding the experimental observations.

To simulate the effects of an ATPase mutation, we abolished the ATP-hydrolyzing activity of one ATPase, for example Rpt3, in the simulation, and then analyzed the global dynamics of the mutated proteasome^29^. ATP hydrolysis at the Rpt3-Rpt4 pocket is important for the transition from the E_D1_ to E_D2_ cryo-EM state; its inactivation mimics a WB mutation and strongly reduces the transition frequency from the E_D1_-to the E_D2_-equivalent conformation in a FEL simulation. This loss of “flow” was compensated by an increase in the transition to an E_C_-like conformation, so that the overall translocation rate was only reduced by 11% (Fig. 6B, S17). Transition networks with an inactivated ATPase also show an expansion of the E_D_-like conformations in which the ATPases at the−2 or−3 positions are flanked by open interfaces (0=mutant, negative=counterclockwise); this trend is consistent with and may explain a previous cryo-EM analysis on WB mutant proteasomes^11^.

**Figure 6.**
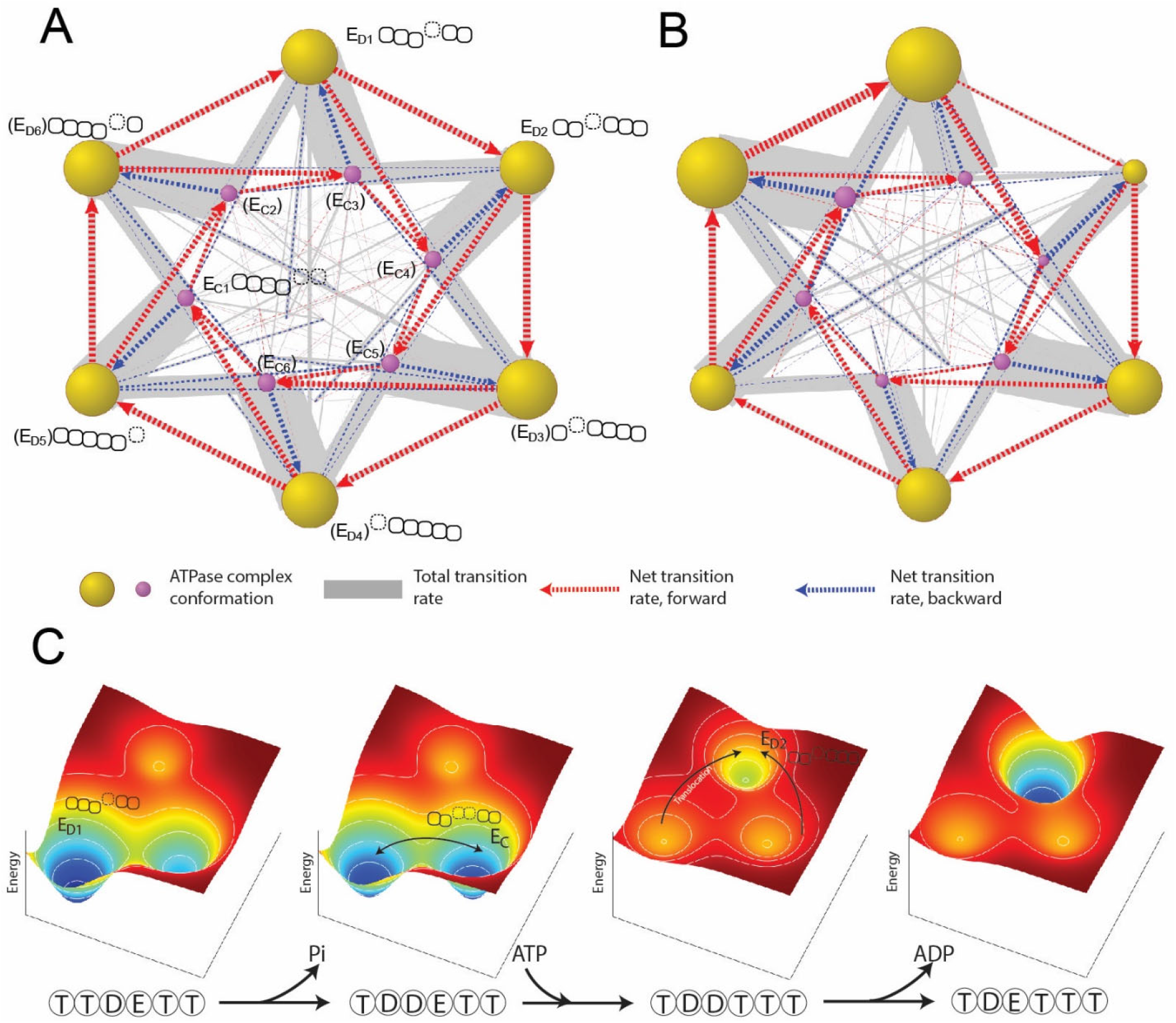
Organization of proteasomal conformations in the dynamical space. **A.** A diagram showing the steady-state transitions between the ATPase complex conformations in the FEL model. Each node represents a unique hexamer conformation, whose size is in proportion to its probability of occupancy in a steady state. Yellow and purple nodes are the E_D_-like and E_C_-like conformations respectively, with the corresponding ATPase architecture as presented in Fig. 2C. The labels of the conformations that have not yet been experimentally identified are bracketed. The width of a grey line is proportional to the total rate of forward and backward transitions between two conformations; and a dashed arrow represents the net transition rate and is colored according to the direction of substrate translocation associated with this conformational change. Minor conformations are randomly distributed in the graph. Table S1 contains a full list of the conformations and their transition rates. The transition diagram without the hydrolysis activity of Rpt3 is shown in **B. C.** The mechanism of the cooperative movements of proteasomal ATPases during substrate translocation. The FELs on three conformations (the E_D1_- and E_D2_-equivalent and an E_C_) are illustrated for four nucleotide statuses in a typical E_D1_-to-E_D2_ transition (Fig. S18). The ATPase architecture of each conformation is presented as in Fig. 2C. Arrows indicate the major conformation transitions upon the change of the nucleotide status and the associated FEL.

The order of the elementary steps of reactions and transitions is an important part of the ATPase mechanism and is challenging for direct experimental study. Different sequences of these steps have been hypothesized to drive the orderly conformational transitions of the proteasome and other AAA+ ATPases^7,18,30^. Our simulation identifies the most-likely reaction sequence by enumerating all the possible sequences that can promote transitions between two states and calculating their probabilities. For the E_D1_-to-E_D2_ transition, the most-likely reaction sequence begins with ATP hydrolysis at the Rpt3-Rpt4 pocket, followed by ATP binding to Rpt5 and an energy-dissipating conformational change. This conformational change exchanges the ADP-bound Rpt3-Rpt4 interface for an ATP-bound Rpt5-Rpt1 interface and drives a unit-step translocation of two AAs by rearranging the PL1 staircase (Fig. S18). This change also drives Rpt4 into a disengaged APO state with a weak nucleotide affinity, allowing rapid release of bound ADP. Otherwise, the rate of ADP release would be incompatible with the fast kinetics of substrate translocation. Phosphate release is unlikely to be limiting since phosphate has a *K_i_* values of 70mM measured in a competition assay, a much weaker affinity than ADP for the ATPases (Fig. S3). Identifying this typical reaction sequence is critical for obtaining the formula for the EC_50_ value of ATP (Methods 10). In a proteasome with non-hydrolyzing Rpt3, the E_D1_-to-E_C_ transition that partially rescues the “flow” is initiated by ATP hydrolysis at the Rpt6-Rpt3 pocket. The associated conformational change drives a translocation of two unit-steps, or four AAs (Fig. S19). These larger steps occur stochastically even in the wildtype proteasome, though less energetically favorable. Nonetheless, they likely explain the experimentally-observed translocation efficiency of 2.6~3.0 AAs per ATP consumption (Fig. 5A)^26^. Variations in stepping size have been observed in substrate translocation by ClpXP, though its connection with the simulation result is unclear^25,31^.

Although we assumed identical parameters for the six ATPases in the FEL model, the model can be extended to interpret the observed functional disparity in proteasomal ATPases. We hypothesize that at least part of the disparity may be caused by the proteasome Lid-ATPase interaction, which breaks the symmetry at the level of hexamer conformations. The Lid subcomplex interacts extensively with the ATPase domains, primarily on Rpt3/6, in the E_A_ or E_A_-like states in which the PL1s on Rpt3 and Rpt2 respectively occupy the top and bottom niches in the staircase and Rpt6 is in a disengaged position^7,32–34^. Such an arrangement closely resembles one E_D_-like conformation (Fig. S20, grey node). Weakening the Lid-ATPase interaction by mutations reduces the fraction of proteasome in the E_A_-like states^35^. We therefore hypothesize that the Lid-ATPase interaction may effectively stabilize this specific ATPase conformation by lowering its standard free energy. We implemented such a stabilizing effect in the FEL simulation by lowering the free energy of this E_D_-like conformation by an arbitrary value, and recorded significantly-expanded occupancy in the E_D1_- and E_D2_-conformations and a simultaneous shrinkage in other E_D_-like conformations (Fig. S20), consistent with the cryo-EM observations^7,8^.

To study whether the Lid-ATPase interaction may contribute to the different growth phenotypes in yeast WB mutants, we examined how the translocation rate changed after simulated inactivation of the ATP-hydrolysis activity of individual ATPases. We found that the predicted translocation rate was qualitatively correlated with the corresponding growth phenotype in a range of the Lid-ATPase interaction strength, which suggests that this interaction may contribute to the growth phenotypes of yeast WB mutants, potentially in conjunction with other compensatory mechanisms (Fig. S21).

In summary, simulations based on the empirical FEL reconstruct *in silico* the dynamics of the proteasomal ATPase complex and provide mechanistic insights into the observed disparities between the individual ATPases in previous structural and functional studies. The steady-state distribution of ATPase conformations is consistent with the cryo-EM observations in multiple studies. These results further support the validity of the FEL approach.

## Discussion

Recent high-resolution cryo-EM studies provide snapshots of the molecular details of the AAA+-family ATPases including the proteasome. Revealing the exquisite dynamics of these complex protein machines remains challenging for both experimental investigation and molecular simulation.

The FEL is an intrinsic property of a protein and is a key component in physical models which have provided important insights into the mechanisms of molecular motors^36–38^. FEL can be derived from conformational occupancies in cryo-EM studies through the Boltzmann equation^39,40^. This approach may not be directly applied to complex and active ATPases such as the proteasome. In this work, we exercised the principle of parsimony to reconstruct the FEL of the proteasomal ATPase complex and experimentally determined its parameters in an attempt to uncover the mechanism in polypeptide translocation. The FEL-based simulation generates various predictions to independently test the simulated dynamics of proteasome and this overall approach. We found that the simulation recapitulated a number of important experimental observations including degradation kinetics, state distributions in cryo-EM datasets and the growth phenotypes of WB mutants (Fig. S7). Moreover, this study reveals that these varieties of phenomena are in fact driven by a coherent and much-simpler principle, thus offering mechanistic insights into the ATP-driven cooperative actions of the ATPases in promoting the translocation of substrate polypeptides.

The observation that the nucleotide status of three consecutive ATPase subunits on the proteasome represents a continuous sequence in an ATP cycle has led to a sequential rotary model for proteasomal ATPase activity^2,7,8^. A strict sequential mechanism requires that the ATP-bound subunit adjacent to the ADP subunit must hydrolyze ATP first. We compared the nucleotide-interacting motifs in all high-resolution structures of the proteasome, but failed to identify a consistent trend that could suggest an allosteric effect between the ATP-pocket and other parts of the complex. As an alternative, we considered the possibility that the apparent cooperativity between different subunits might emerge from the modulation of their collective FEL by ligand binding, as proposed in the celebrated Monod-Wyman-Changeux concerted model for allosteric regulation^41^. The inter-subunit signaling motif has been suggested to mediate ATPase communication and coupling with nucleotides and may underlie the *E_Br_* in the FEL model^42^. Further studies should reveal the molecular basis of how these ATPases communicate.

The ATPase heterohexamer contains at least two open interfaces in all the cryo-EM structures except for E_A_ states, where the “disengaged” subunit is associated with one open interface. This is perhaps due to the extensive Lid-ATPase interaction in these states^32^. The presence of an empty nucleotide pocket entails a unique energy minimum in the FEL and stabilizes the conformation of the ATPase complex at a given nucleotide status. Changing the nucleotide status alters the FEL, drives the conformation of the ATPase complex towards the new energy minimum, and elicits coordinated movements of the individual ATPases (Fig. 6C). Loss of ATPase cooperativity due to flattening of the FEL at above-physiological ATP concentrations reduces the rate of substrate degradation (Fig. 4B). A similar inhibitory effect has been reported for a bacterial pilus assembly motor, though the exact mechanism is still unclear^43^. Although sacrificing some translocation efficiency, the flat FEL of the all-ATP state facilitates interchange among proteasomal conformations, even under physiological ATP concentrations. This property may be important for bypassing an occasional stuck or defective ATPase subunit that would completely inhibit proteasomal activity in a strict sequential-transition model. The resulting network of conformational transitions is analogous to the “ring-resetting” model for the ClpXP ATPase complex^44^. The peptidase activity of proteasome is also inversely correlated with ATP concentrations above a threshold; however, this is likely due to a different mechanism since the critical concentration is ~50x lower than that for translocation^45^.

The nonequilibrium transitions of the ATPase complex identified by the FEL approach reveal important features of proteasome functions. Our model shows that substrate translocation steps are directly coupled to an energy-dissipating conformational transition which swaps an ADP-bound closed interface with another ATP closed interface (Fig. S18). We propose that this coupling may dictate the directionality of translocation. Partial degradation, or processing, by the proteasome is a natural process in the maturation of certain protein factors^46^. The backward process is much less understood, and may contribute to efficient release of partially degraded substrates. In this study, the mechanical force on a substrate during translocation was simplified as a constant. Incorporating the precise unfolding landscape of a substrate and the interaction energy with the ATPase into the FEL model would be straightforward^47^, and may shed light on the mechanism of partial or nonprocessive degradation of certain substrates.

An equivalent mutation on the six ATPases may have different effects on proteasome activities^10,11,13^. The extent to which the structural divergence among the six ATPases contributes to these functional heterogeneities is unclear. Interestingly, we found that interpreting at least some of these functional heterogeneities do not require assuming disparity at the level of individual ATPases, raising the question how these heterogeneities originate. Remarkably, the introduction of Lid-ATPase interactions as a simple conformation-stabilizing parameter in the FEL simulation, recapitulates the asymmetric cryo-EM state distributions and may explain the different phenotypes of WB mutants. Direct interpretation of WA mutants remains challenging, as these mutations often lead to proteasome assembly defects^13^. Predictions by the FEL model that includes the Lid-ATPase interaction are compatible with all the kinetic measurements in our study, with the exception that the peak translocation rate is underestimated by 20%-25%, suggesting incomplete understanding of the symmetry-breaking mechanism.

Protein substrates and ubiquitin chains activate the ATPase activity of the proteasome by 2~4 fold, larger than the predicted increase by the FEL model^48^. This activation process may be independent of the processive translocation phase described by the FEL model, and relate to a transition from the resting state to translocating states of the proteasome^7,49^. Similarly, substrates whose degradation is limited by steps other than translocation, such as the DHFR-containing substrates in this study, may exhibit different characteristics in kinetic studies; examining these substrates may allow the FEL model to be extended to understand other important aspects of the proteasome’s activities.

The FEL model is built on several empirical rules that are generalized from available structures. The FEL presented here is undoubtedly an approximation of the actual FEL of the proteasome, but nevertheless is sufficient to explain several of the key features of the ATPase complex. Other aspects of the structural information, such as the conformational occupancy and nucleotide distribution, are not required for model construction but are still consistent with its predictions (Fig. S7). Identification of additional allosteric effects and minority states of the proteasome should lead to a more precise FEL and further insights into its biological activities.

We expect this FEL approach to have many applications in guiding experimental design and data analysis, and to provide valuable insights into the mechanism of proteasomal ATPases and other AAA machines.

## Acknowledgements

We thank Huang L. for sharing constructs for proteasome purification and thank Schulman B. and Brown N. for sharing constructs for APC/C purification. We are grateful for the critical reading and comments by M. Kirschner, Y. Tu, T. Mitchison, D. Finley, A. Goldberg, L. Bai, R. Ward, J. Yan, and P. Ho. Portions of this research were conducted on the O2 High Performance Compute Cluster, supported by the Research Computing Group, at Harvard Medical School.

## Authors contributions

Y. L. designed the experiments; Y. L., R. F., and J. H. performed the experiments and data analysis; M. Z. analyzed the cryo-EM dataset; All participated in manuscript preparation.

## Data and materials availability

All data is available in the main text or the supplementary materials. The original data is available upon request. Experimental materials are available upon request.

## Supplementary Materials

Materials and Methods

Figures S1-S21

Table S1

References

